# A genome-wide association study of neonatal metabolites

**DOI:** 10.1101/2023.11.25.568687

**Authors:** Quanze He, Hankui Liu, Lu Lu, Qin Zhang, Qi Wang, Benjing Wang, Xiaojuan Wu, Liping Guan, Jun Mao, Ying Xue, Chunhua Zhang, Yuxing He, Xiangwen Peng, Huanhuan Peng, Kangrong Zhao, Hong Li, Xin Jin, Lijian Zhao, Jianguo Zhang, Ting wang

**Author notes:** These authors contributed equally (Quanze He, Hankui Liu). Correspondence: Prof. Ting Wang, The Affiliated Suzhou Hospital of Nanjing Medical University, Suzhou 215000, Jiangsu Province, China;. Prof. Jianguo Zhang, BGI-Shenzhen, Shenzhen 518083, China;. Prof. Lijian Zhao, BGI Genomics, BGI-Shenzhen, Shenzhen 518083, China;. Prof. Xin Jin, BGI-Shenzhen, Shenzhen 518083, China;.

## Abstract

The hereditary component significantly influences the concentration of metabolites in adults. Nevertheless, the precise influence of genetic factors on neonatal metabolites remains uncertain. To bridge this gap, we employed genotype imputation techniques on large-scale low-pass genome data obtained from non-invasive prenatal testing. Subsequently, we conducted association studies on a total of 75 metabolic components in neonates. The study identified a total 17 previous reported associations and 13 novel discovered associations between single nucleotide polymorphisms and metabolic components. These associations were initially found in the discovery cohort (8,744 participants) and subsequently confirmed in a replication cohort (19,041 participants). The average heritability of metabolic components was calculated to be 76.2%, with a range of 69-78.8%. The aforementioned findings offer valuable insights pertaining to the genetic architecture of neonatal metabolism.

**In Brief:** Large-scale genomes of maternal non-invasive prenatal testing provide insights into the genetic contribution to neonatal metabolism.

**Highlights:** GWAS of 27,785 low-pass genomes revealed 13 novel associations of neonatal metabolic components.

Estimated an average of 76.2% heritability of neonatal metabolic components and showed the individual concentration can be accurately predicted from polygenic risk scores.

A total of 17 established relationships have been observed, providing evidence that maternal genomes can be utilized in neonatal metabolite GWAS.

## Introduction

Metabolites play an important role in biological function in newborns.^1,2^ Abnormal concentrations of metabolites are considered to be harmful factors in the context of inherited metabolic disorders. In recent years, a multitude of genome-wide association studies (GWAS) have identified hundreds of loci related to metabolites in the adult population. These findings provided compelling evidence for the role of genetic factors in contributing to the variations of metabolite concentrations.^3–8^ However, the genetic contribution to neonatal metabolism remains largely unexplored. Furthermore, metabolism serves as a link between genetic factors and human traits. Lipid-related genes were shown to involve healthy infant growth.^9,10^ Inherited metabolic illnesses, such as propionic acidemia^11^ mucopolysaccharidoses,^12^ and tyrosine hydroxylase deficiency^13^ have been found to be caused by severe genetic mutations. The examination of the genetic component in neonatal metabolism requires attention.

Currently, the routine measurement of DNA in newborn dried blood spots is not commonly practiced,^14^ posing a significant obstacle for the genetic investigation of neonatal metabolism. The goal of accessing low-pass whole-genome sequencing (WGS) data of non-invasive prenatal testing (NIPT) can now be reliably achieved with recent technological advancements in genotype imputation. Previous studies^15–17^ have demonstrated that large-scale low-pass WGS can be used for GWAS analysis and polygenic risk score (PRS) calculation. In our previous study, we employed genotype imputation techniques on a dataset consisting of 141,431 low-pass genomes of cell free DNA (cfDNA). This analysis yielded noteworthy findings regarding the genetic associations of height, body mass index (BMI), and twin pregnancy, which were previously unknown.^18^ By employing the GWAS methodology, which evaluates the direct and/or indirect association between genetic locus and phenotype,^19^ it becomes feasible to link the WGS data of the mother and the phenotypes of newborn metabolic screening.

In this study, we performed genotype imputation on 27,785 maternal low-pass genomes of cfDNA and conducted a two-stage GWAS on 75 metabolic components in neonates. These components included 43 independent metabolites and 32 ratios. We have identified a total of 17 previously reported loci and 13 newly discovered loci that are associated with metabolic components. Additionally, we have proposed a robust approach for conducting GWAS on neonatal metabolism by using maternal genotypes as a link.

## Results

### Participant characteristics, sequencing features, variant calling, and genotype imputation

In our study, a total of 27,785 WGS datasets of maternal NIPT were chosen, covering the period from 2015 to 2020. The majority of pregnant women originated from Jiangsu Province, accounting for 60% of the total. The distribution of maternal ancestry in different regions of China is as follows: 14% in north China, 79% in central China, and 7% in south China. A qualification analysis of 43 independent metabolites, including 11 amino acids, 31 carnitines, and succinylacetone, was conducted on neonatal dried blood spots obtained at day 3 after delivery. Additionally, 32 ratios were derived based on the measured metabolites (Table S1; Figure S1). By considering the systematic bias of sequencing platforms,^18^ we designed a two-stage GWAS and integrated the statistical summaries for a meta-analysis. A total of 27,785 paired data were classified into two distinct cohorts according to the sequencing platform utilized for NIPT. Specifically, the discovery cohort consisted of 8,744 genomes that were sequenced by Ion Torrent, while the replication cohort comprised 19,041 genomes that were sequenced by Illumina CN500.

During the stage of discovery, pregnant women received low-pass WGS with an average of five million single-end reads (150bp). The reads were aligned to the hg38 reference by bwa software, resulting in an average sequencing depth of 0.28× and coverage of 25% of the genomic region (Figure S2A). During the stage of replication, it was observed that an average of five million short single-end reads (35bp) resulted in an average sequencing depth of 0.06 and coverage of 5% of the genomic region (Figure S2A). Via the utilization of BaseVar,^18^ a total of 58 million single-nucleotide variants (SNVs) were identified and the mutant allele frequencies (MuAFs) were estimated. SNVs that exhibited low quality or abnormal depth (Figure S2B) were excluded from the analysis. A total of 21 million single nucleotide polymorphisms (SNPs) were selected for genotype imputation, based on their minor allele frequency (MiAF) > 1% and a transition-to-transversion ratio (Ti/Tv) of 2.12. The imputation process was performed by STITCH^20^ utilizing a reference panel consisting of Han Chinese individuals from the 1000 Genomes Project. A set of 5.4 million well-imputed SNPs were reserved based on specific criteria: MiAF > 1%, *P*-value of Hardy-Weinberg balance test > 1×10^-5^, imputation accuracy (I_A_) > 0.4, and genotyped rate > 0.9. Most of the SNPs (91.7-99.9%) with MiAF > 5% were found in the dbSNP, ChinaMAP, and/or gnomAD-EAS databases. Additionally, a percentage (26.1-36.9%) of low-frequency SNPs with MiAF < 5% were discovered to be novel (Figure 1A). To estimate the accuracy of imputation, a comparison was made between the MuAFs calculated by STITCH and another three sources of MuAFs, namely the ChinaMAP database, gnomAD-EAS database, and BaseVar method. A high density of SNPs was seen along the diagonal line (y = x), accompanied by statistically significant correlations (R^2^ ≥ 0.99, β = 1) of MuAFs for these SNPs (Figure 1B). These findings provided evidence that the MuAFs estimated by STITCH can be considered trustworthy. In the process of replication, a total of 19,041 cram files were employed to impute the individual genotypes of the 21 million SNPs that were identified during the discovery phase. By applying identical filter criteria for the purpose of identification, we successfully preserved a total of 1.5 million well-imputed SNPs. Furthermore, we observed a robust MuAF consistency (R^2^ = 0.99, β = 0.99) between the initial discovery and subsequent replication stages (Figure 1B).

**Figure 1.**
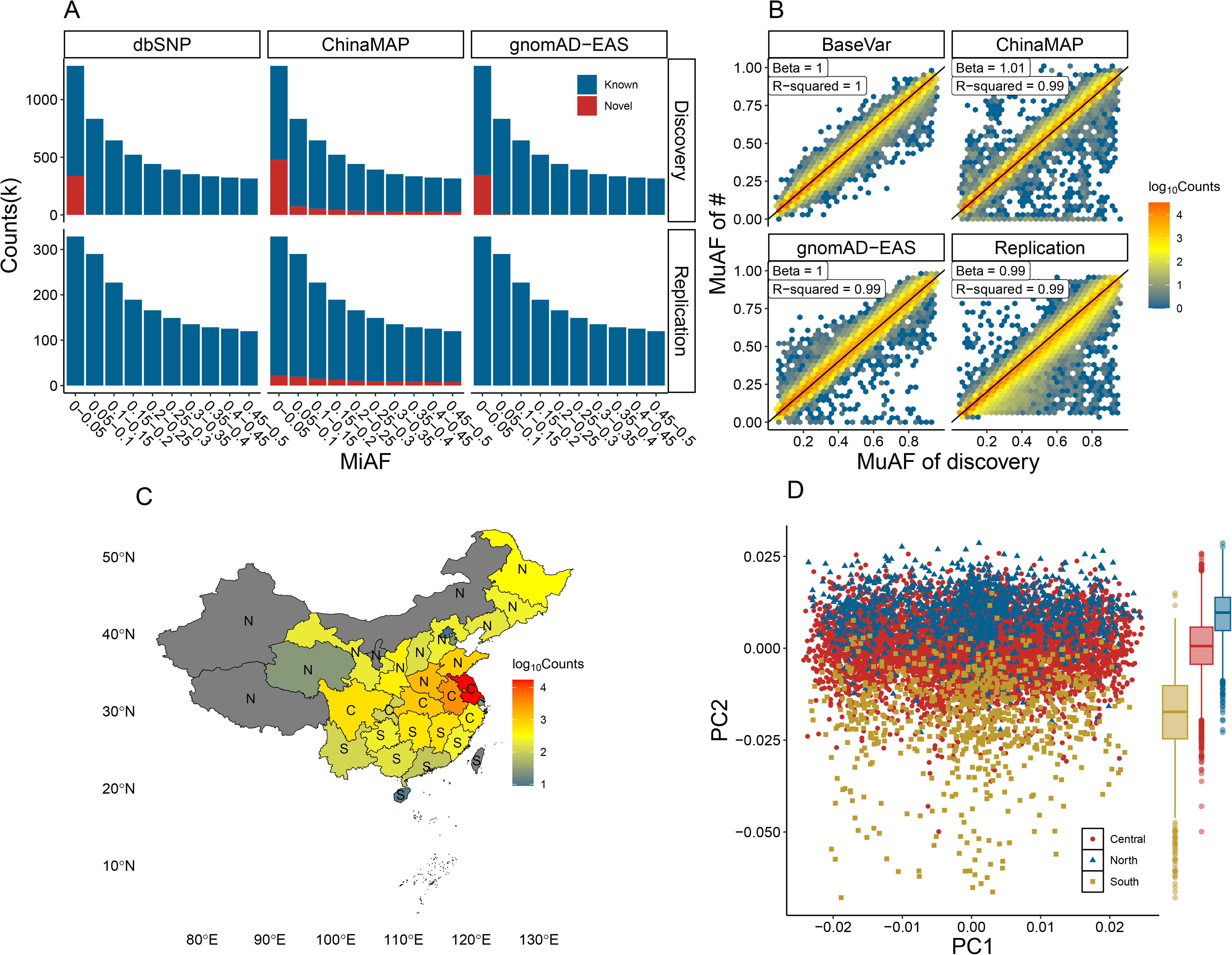
Allele frequency spectrum and genetic structure. Comparison of known and novel SNPs in dbSNP, ChinaMAP, and gnomAD-EAS database (A). The accuracy of allele frequency of well-imputed SNPs (B). The geographic distribution of participants (C) and their genetic structure (D).

Utilizing the well-imputed SNPs, we conducted a principal component analysis (PCA) to discern distinct genetic components among individuals from the northern, central, and southern regions of China (Figure 1C). This analysis revealed that genetic component exhibited by the participants align with their respective geographical origins (Figure 1D). This finding consisted to our previous study^18^ which employed genotype data identified by BaseVar for PCA. Additionally, we noted a correlation between the second principal component (PC2) and fetal cfDNA, which proportion was estimated from 1-30% with a median of 10% and affected by the maternal BMI (Figure S3A). This finding indicated that fetal cfDNA has an impact on individual genotypes. It is imperative to establish a correlation between the proportion of cfDNA and neonatal metabolism. Linear regression analysis did not yield any statistically evidence of a significant association (Figure S3B), indicating that there is no observable bias in the individual genotypes affected by the fetal cfDNA for GWAS of neonatal metabolism. On the other hand, maternal age/height/BMI and neonatal weight/sex were found to strongly influence the neonatal metabolism. These factors were utilized as covariates in later GWAS (Figure S3B).

### Genome-wide association of maternal height/BMI and neonatal metabolic components

We performed a GWAS to investigate the association between maternal height and BMI. Three SNPs that are linked to maternal height or BMI were found in the discovery phase and then confirmed in the replication phase (Table S2). Two SNPs (rs7571816, rs545608) were documented to associate with the consistent phenotype in the GWAS catalog.^21^ *DIS3L2* (rs7571816) has been found to be significantly associated with body height.^22,23^ *CRYZL2P-SEC16B* (rs545608) has been found to have associations with BMI,^24^ Type 2 diabetes,^25^ and body fat.^26^ The rs7206410 is located in *FTO* gene, which has been linked to obesity.^27^ In our previous study,^18^ we also observed an association between rs7206410 and BMI. Collectively, the established associations substantiate the reliability of our study’s design and findings.

In the present study, we established an association between maternal genetic variants and their neonatal metabolic components in order to reveal the involvement of genes related to metabolism. The genomic control factors (λ) for 72 metabolic components were found to < 1.1, except for C6DC, C10:1, and C12:1 (Figure S4). This suggests that the linear regression model, which included neonatal weight/gender, maternal age/height/BMI, and the top three PCs as covariates, effectively adjusted for genetic inflation. A total of 2,628 associations between SNPs and metabolic components were detected by a significant threshold of *P*-value < 5×10^-7^. Out of these associations, 1,863 were seen in the replication cohort. Furthermore, 1,162 associations were successfully replicated using a threshold of *P*-value < 2.68 × 10^-5^ (adjusted for multiple comparisons at a significant level of 0.05 based on the 1,863 associations in the replication cohort). In the meta-analysis, a total of 1,104 associations, consisting of 414 SNPs, were certified as statistically significant based on a threshold of *P*-value < 6.67 × 10^-10^. This threshold was determined by correcting the genome-wide significant threshold of 5 × 10^-8^ by taking into account 75 components (Figure 2). In order to determine the tag SNP, we selected the SNP exhibiting the highest level of statistical significance, as shown by the *P*-value, inside a gene or a region spanning one million bases that lies between genes. Subsequently, we kept a total of 30 associations pertaining to 13 neonatal metabolic components (Table 1).

**Figure 2.**
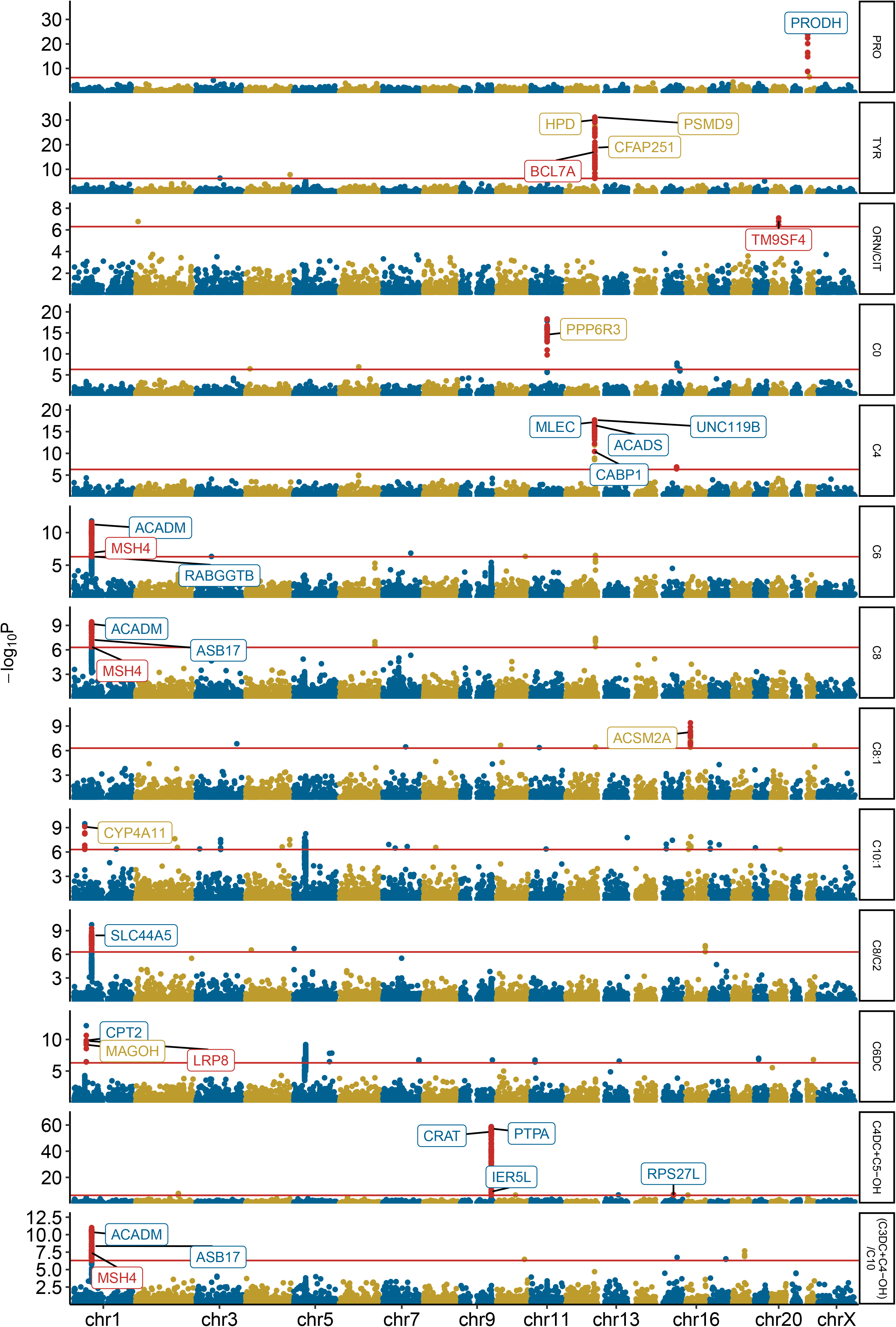
Manhattan plots of 13 metabolic components. SNPs met all of the thresholds in discovery (< 5 × 10^-7^), replication (< 2.68 × 10^-5^), and meta-analysis (< 6.67 × 10^-10^) are indicated by red points. Genes with blue color indicate the 14 genes of 17 known associations documented in GWAS catalog. Genes with yellow color indicate the 7 genes of 7 novel associations that the SNPs were documented to be associated with other metabolic components. Genes with red color indicate the 4 genes of six novel associations. Y-axis refers to the *P*-values of GWAS in discovery. Red horizontal line refers to the significant threshold (5 × 10^-7^) in discovery.

**Table 1.**
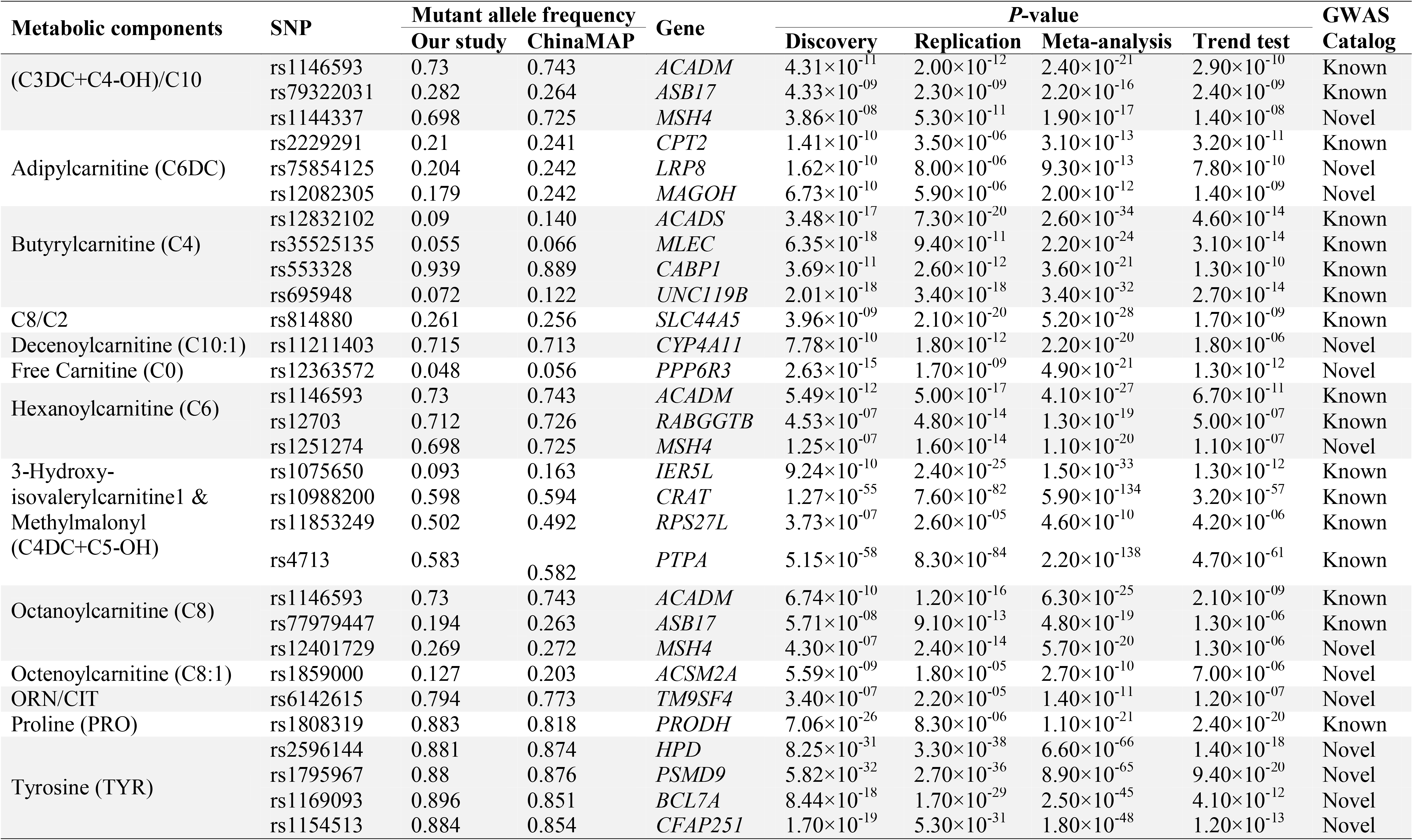

Here, we have linked maternal SNPs and neonatal metabolism. In order to establish the genetic links between metabolism and the neonatal population, we employed a trend test^28^ to test the associations between the frequency of neonatal alleles and the ranking of phenotypic groups. The quantitative metabolic components were ranked and the neonatal participants were subsequently categorized into five distinct groups based on these rankings (Figure 3A). In accordance with Hardy-Weinberg balance, the frequency of allele in the offspring population is equivalent to the frequency of allele in the parental population. The SNPs associated with metabolism successfully passed the Hardy-Weinberg balance test, indicating that they conform to the expected genetic equilibrium. The allele frequency of these SNPs was analyzed in a sample of 602 trios with high-depth WGS data.^29^ It was noted that there was a significant similarity in the allele frequency between the offspring and the parental population (Figure S5A). Furthermore, an analysis of the allele frequencies in the gnomAD-EAS database revealed that these frequencies were comparable between males and females (Table S3). Based on these facts, it can be inferred that the estimation of allele frequency in neonatal groups can be derived from maternal groups. A novel approach (Methods) was devised to determine the neonatal allele frequency based on the maternal allele frequency and gnomAD database, which provides information on allele frequencies in both males and females. The accuracy of these estimations was subsequently validated by the WGS data of 602 trios (Figure S5B). Through the utilization of trend test, we demonstrated a positive correlation between neonatal allele frequencies and the corresponding increase in neonatal metabolic concentration across the five ranked groups (Figure 3B). The findings offer statistical evidence for the validation of the 30 associations between neonatal genotypes and metabolic components.

**Figure 3.**
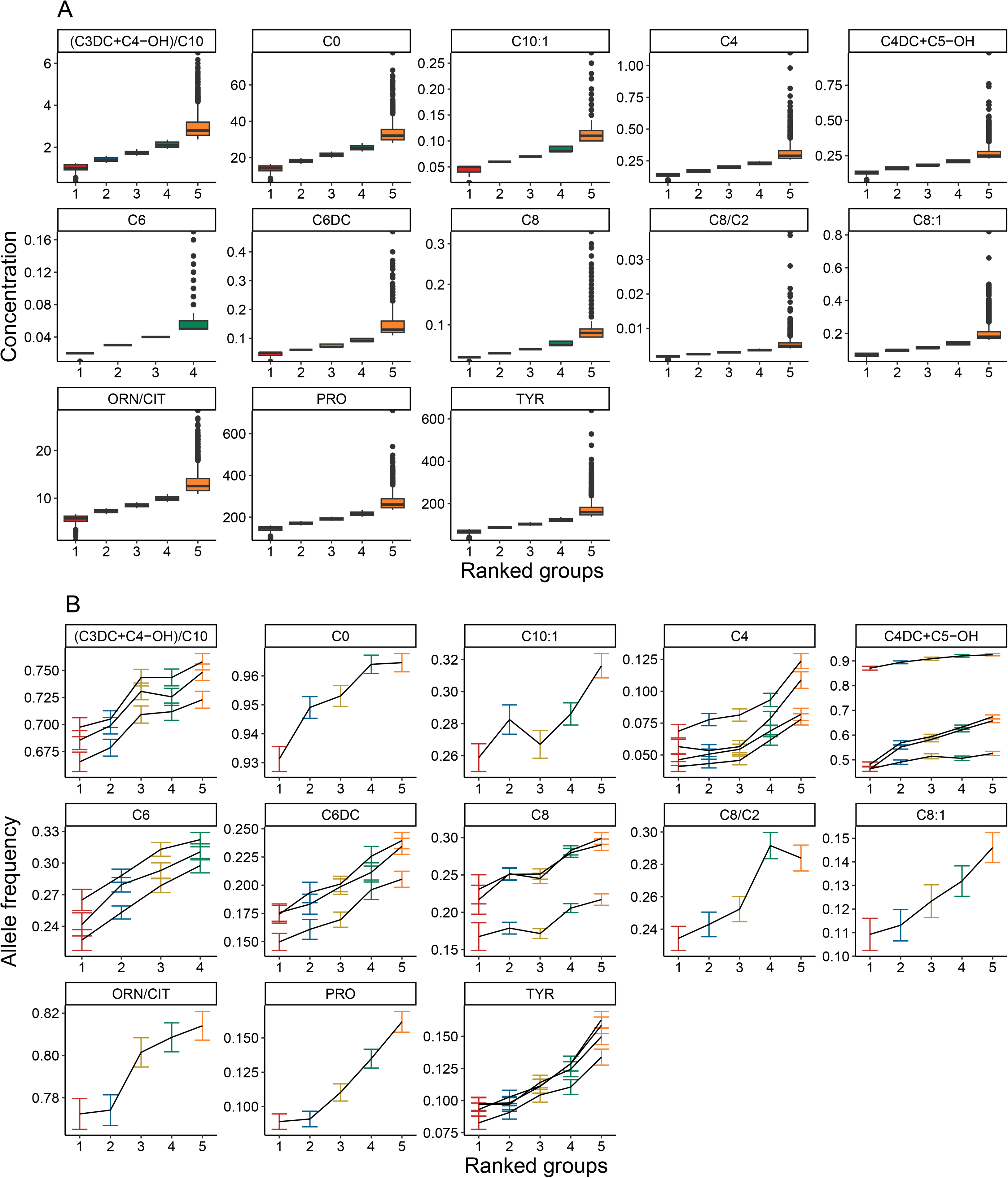
The correlation of allele frequency of SNP and metabolic concentration in neonates. The distribution of each metabolic concentration of five ranked groups (A). Allele frequency of tag SNP increased following the increase mean of metabolic concentration among the five ranked groups (B).

Among the 30 associations, 17 associations (14 genes: *ACADM, ASB17, CPT2, ACADS, MLEC, CABP1, UNC119B, SLC44A5, RABGGTB, IER5L, CRAT, RPS27L, PTPA, PRODH*) were documented in the GWAS catalog. Out of the 13 novel associations examined, seven associations involving the genes *MAGOH, CYP4A11, PPP6R3, ACSM2A, HPD, PSMD9,* and *CFAP251* were found to be associated with various metabolic components (Table 1). On the other hand, six associations (specifically, rs6142615: *TM9SF4*; rs1144337/rs1251274/rs12401729: *MSH4*; rs75854125: *LRP8*; rs1169093: *BCL7A*) have not been previously reported in the GWAS catalog (Figure 4A; Table S3). The involvement of *TM9SF4* in metabolism was reported for the first time. The genes *MSH4, LRP8,* and *BCL7A* are located in close proximity to the already identified loci associated with metabolism. The presence of known SNPs and genes associated with metabolism demonstrated that the maternal NIPT WGS data has potential for investigating the genetic associations for neonatal metabolites.

**Figure 4.**
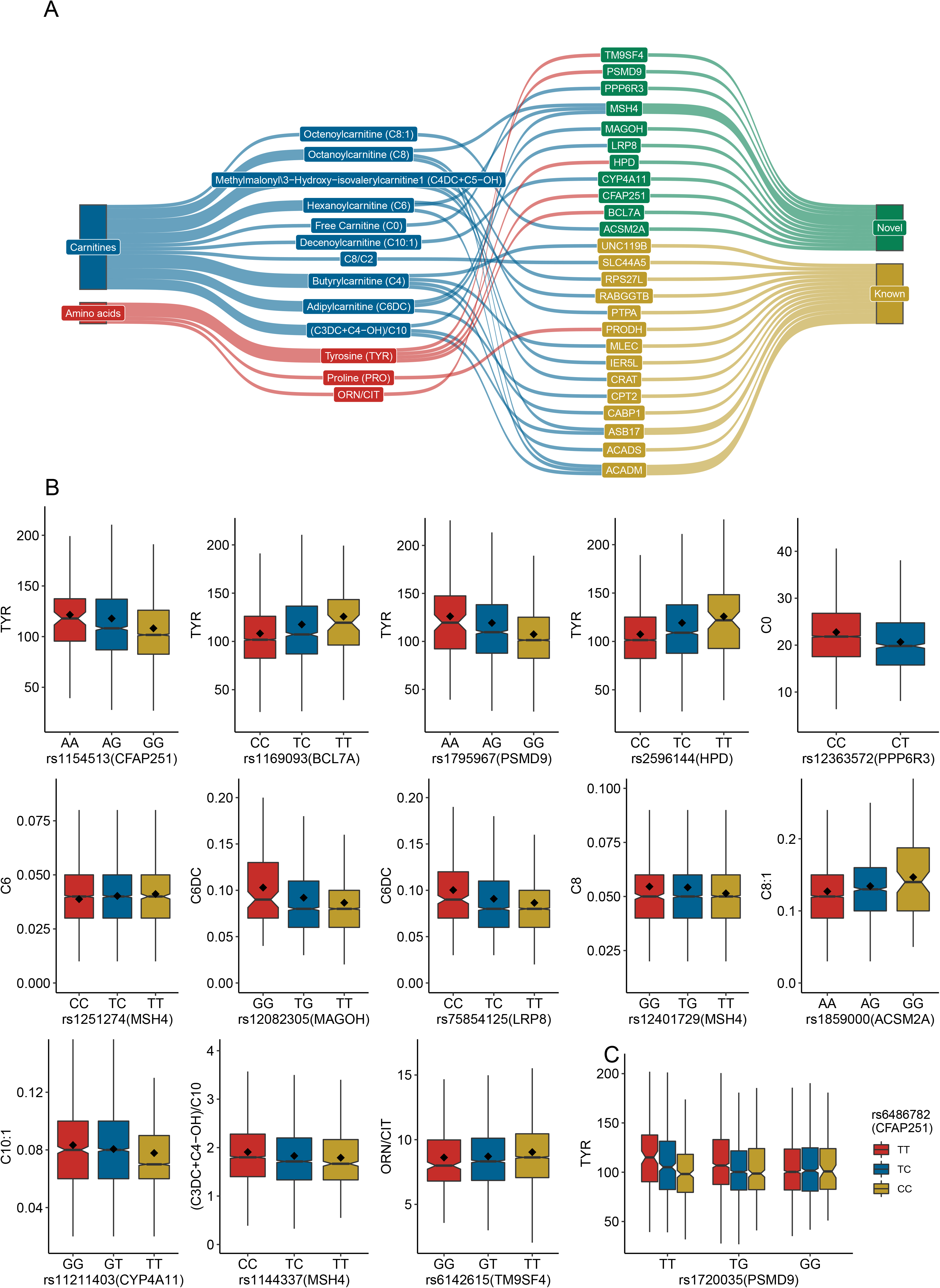
The connections of metabolic components and related genes were displayed in A. The SNP effects of the 13 novel significant associations were displayed in B. A pairwise SNP-SNP interaction was displayed in C.

In the context of the 13 novel associations, our findings demonstrated an additive allele effect of the SNPs in their influence on metabolic concentrations (Figure 4B). The presence of the rs2596144 T allele in the *HPD* gene has been demonstrated to result in elevated tyrosine levels in neonates. The *HPD* gene has been documented as a causative factor in type III tyrosinemia (OMIM:276710).^30^ Pathogenic homozygous mutation in the *HPD* gene results in a significant deficiency in the activity of 4-hydroxyphenylpyruvate dioxygenase. This deficiency is characterized by abnormally high levels of tyrosine in the bloodstream (Table S4). Furthermore, a significant interaction of two SNPs was observed in relation to the influence on tyrosine concentration (Figure 4C). The statistical significance of the interaction between rs1720035 (*PSMD9*) and rs6486782 (*CFAP251*) was observed in both the discovery phase (*P-*value = 5.32 × 10^-7^) and the replication phase (*P-*value = 5.62 × 10^-7^). This suggests that the impact of rs1720035 is influenced by rs6486782, and that the combined effects of these two SNPs are cumulative. These findings have enhanced our comprehension regarding the influence of genetic factors on metabolic concentration and its association with disorders.

### Knowledge annotation of neonatal metabolism-related genes

In order to explore the primary functional attributes of genes reported in our study, we utilized KEGG enrichment analysis. The result indicated that the metabolism-related genes are implicated in the metabolic pathway of amino acids and lipids (Figure 5A). The PhenoScanner^31^ tool detected a total of 26 distinct phenotypes and/or diseases associated with six novel loci (Figure 5B). The *TM9SF4* gene has been implicated in the regulation of myeloid leukocytes, specifically in relation to monocyte count, and the proportion of monocyte among the total white blood cells.^32^ In our work, we observed that this particular gene had an impact on the neonatal metabolic ratio (ORN/CIT). Citrulline (CIT) could regulate the monocyte percentage of white cells and modulate the regulatory T-cells function differently in infantile rats.^33–35^ Ornithine (ORN) has been found to augment the process of autophagy and exert regulatory control on the infection caused by mycobacterium tuberculosis infection.^36^ In order to investigate the specific organs/tissues/cell types in which these genes exhibit activity, we conducted an expression quantitative trait loci analysis. The result indicated that these genes are active in hepatocytes/liver (Figure 5C). Furthermore, an examination of cell-type expression enrichment in single-cell transcriptomes showed that these genes exhibit distinct expression patterns in the hepatocytes (particularly, the hepatocyte subtype 3) of fetal/adult liver and the proximal tubules of fetal kidney (Figure 5D). The hepatocyte subtype 3 is a specific cell type that exhibits a high level of gene expression related to lipid, cholesterol, and biosynthesis synthesis.^37^ The kidney is an organ with significant metabolic activity, wherein renal tubules utilize fatty acid oxidation as a means to produce adenosine triphosphate.^38^ However, there was no observed enrichment in any of the cell types of fetal heart or gut.

**Figure 5.**
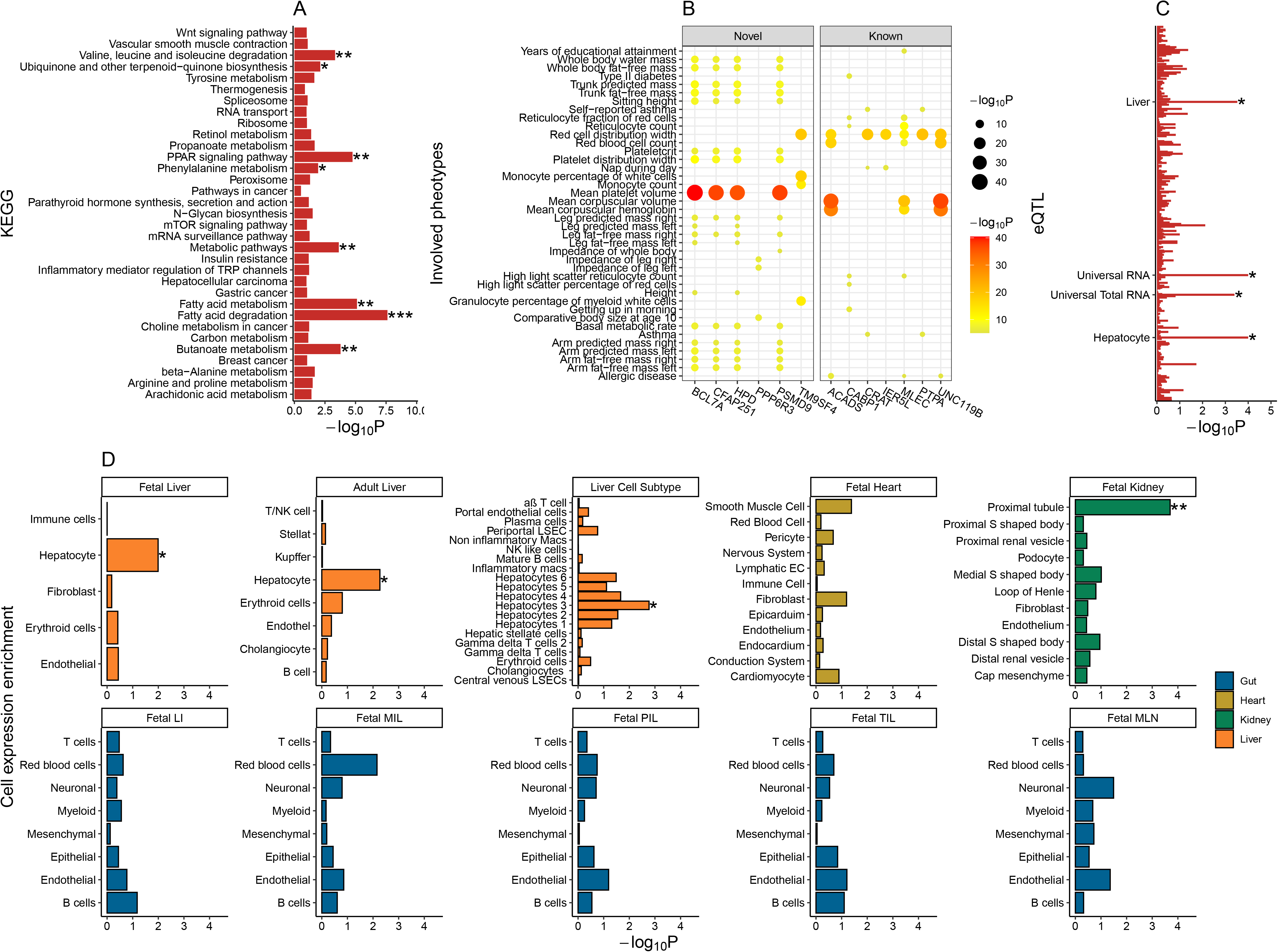
Knowledge annotation of metabolism-related genes. KEGG enrichment analysis annotates eight functional characteristics of the metabolism-related genes (A). PhenoScanner identifies 26 phenotypes related to six novel loci (B). The metabolism-related genes harbor significant enrichments of expression quantitative trait loci in four tissue/organs (C) and specifically expressed in the hepatocytes of fetal/adult liver and proximal tubules of fetal kidney (D). No significant association was observed in any cell types of fetal heart and five regions of fetal gut, including the large intestine (LI), middle ileum (MIL), proximal ileum (PIL), terminal ileum (TIL), mesenteric lymph node (MLN).

### Heritability of neonatal metabolic components

The estimation of heritability using GCTA involves calculating the kinship of individuals based on their genotype matrix, rather than directly using the genotype matrix itself. This approach allows for the estimation of heritability in neonates based on maternal genotypes. In order to validate our hypothesis, we conducted an observation on the WGS data of 602 trios. Our findings revealed a noteworthy association between the pairwise kinship of the maternal population and that of the neonatal population (Figure S5C). The GCTA tool was utilized to analyze the SNPs with a significant level of *P*-value < 0.01 and estimate an average heritability of 76.2% (with a range of 69-78.8%) for neonatal metabolites. Additionally, the heritability estimated for maternal height and maternal BMI were found to be 79% and 80% respectively (Figure 6A). The observed high heritability indicated that genetic factors contribute a major component to the variation of metabolic concentration in neonates.

**Figure 6.**
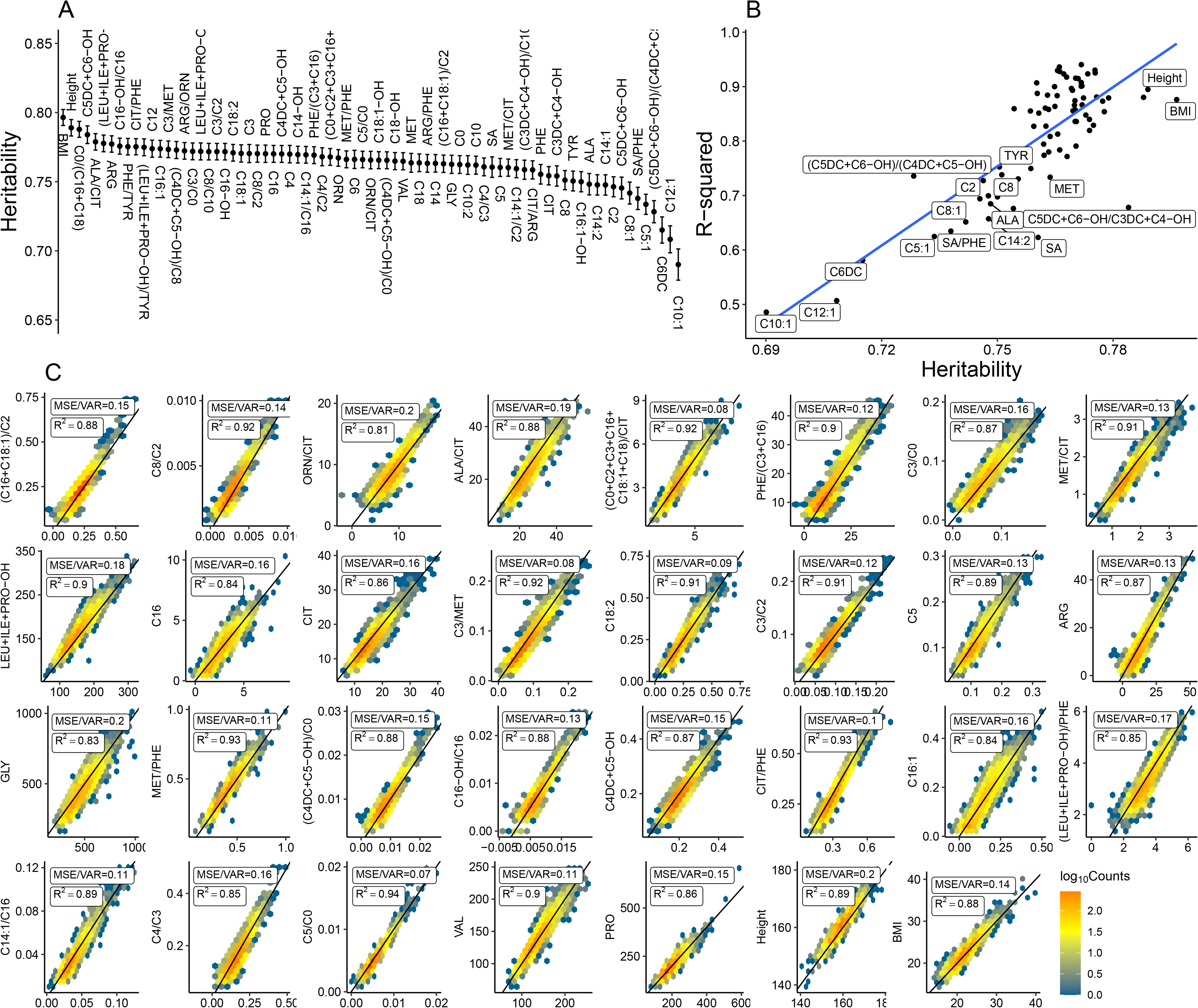
Metabolic heritability and PRS prediction. The black dot and error bar showed the estimated heritability and standard error (A). A correlation was observed between the heritability and R-square of the PRS model (B). The accuracy of prediction was calculated by MSE/VAR. A total of 29 neonatal metabolic components and maternal height/BMI can be accurately predicted by the PRS model using a threshold of MSE/VAR ≤ 0.2. The x and y axis refer to the prediction and observation of concentration of metabolite respectively (C).

The significant heritability of neonatal metabolism suggests the potential to predict the individual concentration of metabolic based on genetic data. Taken together with the addictive effect of associated SNPs, we applied a polygenic risk score (PRS) method to calculate genetic scores for each participant, using the GWAS summary of SNPs with a *P*-value threshold of < 0.01. The results showed a correlation between the R-square of the PRS model and heritability (Figure 6B). The evaluation of prediction accuracy involved the utilization of mean-square error (MSE), which was calculated based on the comparison between the predicted values and the observed values. To facilitate the comparison of accuracy across various metabolic components, the MSE was divided by the variance (VAR) of the observed values. The results exhibited an accurate prediction of 29 metabolic components (MSE/VAR ≤ 0.2), along with maternal height and BMI (Figure 6C). Besides, there exists a strong correlation (R^2^ > 0.8) between observation and prediction for 15 metabolic components. However, it is important to mention that the linear model’s intercept requires adjustments (Figure S6).

## Discussion

Our study carried out a large-scale GWAS on neonatal metabolism. Through this analysis, we were able to identify 17 previously reported associations documented in the GWAS catalog. Additionally, we revealed 13 novel associations, one of which involves a gene (*TM9SF4*) that plays a role in neonatal metabolism of citrulline/ornithine. *TM9SF4* was reported to involve in the osteoporosis and *Tm9sf4*^−/−^ mice exhibited increased bone mass and reduced lipid accumulation in trabecular bones.^39^ Interestingly, citrulline and ornithine are suggested to impact fracture healing.^40,41^ The association of *TM9SF4* and citrulline/ornithine may link a pathway to understand the mechanism of osteoporosis and highlight the gene/metabolite target for the clinical treatment. Moreover, our study demonstrated a significant heritability of neonatal metabolism, together with maternal height and BMI. The findings indicate that metabolic concentrations can be accurately predicted using genetic data, as evidenced by the high heritability and the successful application of the PRS. These results deepen our comprehension of the influence of genetic factors on neonatal metabolism.

Post-GWAS analysis revealed substantial connections between metabolism-related genes and their activation in fetal hepatocytes and kidney’s proximal tubules, indicating the involvement of these two critical cell types in the metabolic processes of newborns. According to current understanding, the liver and kidney are recognized as the primary organs implicated in human metabolism and exert a significant impact on cardiovascular risk within the context of metabolic syndrome.^42^ Hepatocytes are the primary cellular constituents of the liver that plays crucial role in the metabolic processes of carbohydrate, lipid and proteins.^43^ The kidney plays a significant role in the metabolism processes of carbohydrate, with approximate 40% of systemic glucose production occurring in the proximal tubule during periods of fasting and under stress conditions. In contrast, while the intestinal tract does contribute to the metabolic responses^44^ and provides the heart with energy through oxidative metabolism,^45^ there is currently no statistical evidence to support a significant involvement of fetal gut and heart in neonatal metabolism. These findings provide a connection among genetic, cellular, and biological processes, shedding light on the underlying mechanism of neonatal metabolism.

Benefitting by the GCTA^46^ method, we estimated the heritability of neonatal metabolism and maternal height from the maternal kinship. The calculation of heritability is influenced by several factors, including the quantity of significant SNPs and the specific approach employed, such as GCTA^46^ for individual genotype data or LDAK^47^ for statistical summary data. In light of the computational difficulties associated with determining the kinship between pairs participants based on million SNPs, we choose to utilize a subset of SNPs with *P*-value < 0.01 for practicality purposes. While it is possible that heritability may be overestimated when considering a major fraction of significant SNPs rather than all SNPs, our findings indicate a heritability of 79% for maternal height. This aligns with previous estimation of 84-94% in female monozygotic twins and 49-56% in dizygotic twins.^48^ At this particular scale, it is estimated that neonatal metabolism has an average heritability of approximately 76%, yielding a higher heritability compared to the estimation from a sample of 381 newborn twins, consisting of 107 monozygotic and 274 dizygotic twins^49^. The following top 10 components identified in Alul’s study^49^ showed the differences compared with that in our study: C4DC(83% vs. 77%), C4(66% vs. 77%), C5(61% vs. 76%), C2(50% vs. 74.6%), C0(45% vs. 76.2%), C3(44% vs. 77%), C3/C2(64% vs. 77.2%), C4/C3(61% vs. 76.1%), PHE/TYR(51% vs. 77.5%), and TYR(47% vs. 75%). These differences may be attributed to the higher proportion of dizygotic twins, which accounted for 71% of Alul’s study cohort. In adult population, the heritability is approximated to be 50%.^50^ As the phenotypic variance was divided into genetic variance and environmental variance, it was found that the variance in metabolism was largely influenced by genetic factor during the newborn stage, while the influence of environmental factors increased in adult.

In addition to the aforementioned results, our team is currently conducting two supplementary research that independently investigate the genetic influence on maternal metabolites and traits throughout pregnancy. In conjunction with the findings from these two studies, our research showed a correlation between the *HMCN2* gene locus and the concentration of citrulline in both maternal and neonatal subjects. Additionally, we observed an association between the *MARCH8*-*ZFAND4* gene locus and maternal hemoglobin-related traits, as well as neonatal levels of C0 and C2. Furthermore, we identified a relationship between the *SPPL3* gene locus and C-reactive protein levels in mothers, as well as neonatal levels of C4, C4/C3, and C4/C2. These findings suggest a correlation between neonatal and maternal metabolism/traits.

It is important to take into account a potential limitation. The genetic connections in neonates have been demonstrated; nevertheless, the precise biological mechanism underlying the association between maternal genotype and neonatal metabolic concentration remains elusive. While the initial observation of the link was made in relation to the maternal genotype, it is important to note that we did not propose a direct influence of the maternal genotype on neonatal metabolism. In our study, we have put forth a hypothetical mechanism by which the effective allele is transmitted from the mother and/or potentially the unobserved father to the offspring, thereby exerting an influence on the concentration of related metabolite. Our study demonstrated a positive correlation between the allele frequency in maternal/ neonatal groups and the concentration of neonatal metabolite. The obtained outcome provided evidence in favor of our supposition.

The primary inquiry of the study is to the applicability of utilizing genetic data derived from a collection of maternal and fetal cfDNA for GWAS. Fetal cfDNA contribute a proportion ranging from 1% to 30% of the total cfDNA, hence influencing the genotype component. The examination of genotype bias influenced by fetal cfDNA in a NIPT-WGS GWAS necessitates thorough discussion. If there is no observed correlation between the fraction of cfDNA and phenotype, then the imputed genotypes are no biased for GWAS. However, if there is a correlation, it is recommended to include either the proportion of cfDNA and principal components (PC) as covariates in the association model. In our study, no significant evidence was found to support a link between the fraction of cfDNA and neonatal metabolism. The presence of fetal cfDNA during gestational weeks when NIPT is conducted does not appear to have an impact on neonatal metabolism after delivery. This suggests that the proportion of cfDNA dose not introduce bias to GWAS. However, a significant association was seen between the fraction of cfDNA and maternal BMI. The previous work has reported on the association between fetal cfDNA and maternal BMI,^54^ indicating that the inclusion of proportion and/or PC as covariates is necessary in the association test of maternal BMI. Besides, in our study focused on GWAS of neonatal phenotype, we showed the allele frequency of SNPs in Hardy-Weinberg balance is congruent between neonates and their mothers. Furthermore, we have devised a trend test based on allele frequency to validate the association established by maternal genotypes. The findings of this study indicate that a total of 17 known associations, 7 associations documented to be associated with various metabolic components, and 5 associations involving genes neighboring known loci, support the feasibility of this approach in neonatal metabolism GWAS. The widespread global application of NIPT technology, together with subsequent newborn testing, has rendered the utilization of maternal genotype as a means to conduct GWAS on neonatal normal traits highly valuable.

## Methods and Materials

### Study subjects

According to the maternal ID of NIPT and neonatal ID of newborn metabolic screening between 2015 and 2020, 27,785 pairs of maternal NIPT low-pass WGS data and neonatal metabolic data were selected. Participants with positive results on chromosomal abnormalities, abnormal ultrasound fetuses, or abnormal neonatal metabolic screening result have been excluded. 8,744 participants were sequenced by the Ion Torrent platform and 19,041 participants were sequenced by the Illumina CN500 platform. Plasma was divided from maternal peripheral blood (5 mL) by centrifuging and then 600 μL (Ion Torrent) or 1.4 mL (Illumina CN500) plasma was used to extract cell-free DNA. The library construction, quality control, pooling, and sequencing were performed according to their respective sequencing experiments instruction of the Ion Torrent platform and Illumina CN500 platform, respectively. Neonatal metabolites (11 kinds of amino acid, 31 kinds of carnitine, and succinylacetone) were quantified by mass spectrum (Waters TQ Detector) in neonatal dried blood spots.

The written informed consent was approved by Institutional Review Board of Suzhou Municipal Hospital at 2015 and obtained from each participant from 2015-2020. The study was approved by the Institutional Review Board of Suzhou Municipal Hospital (K-2021-024-H01). The genetic/metabolic data collecting are approved ([2022] CJ2133) by the Human Genetic Resources Administration of China.

### Variant calling

Raw sequencing reads with a proportion of gap (N) or low-quality base (Q < 20) at a threshold of > 0.1 were filtered by SOAPnuke.^55^ Clean reads were aligned to human genome reference (GRCh38) by bwa.^56^ Duplicate reads probably generated from PCR were excluded by sambamba.^57^ Reads around known insertions and/or deletions were re-aligned by the indel realignment modules of GATK.^58^ Read qualities were rebuilt by base quality recalibration modules of GATK with references of variants from the HapMap project,^59^ 1000 Genomes project,^60^ and dbSNP database.^61^ Recalibrated reads were stored in cram format by samtools.^62^ The fetal cfDNA proportion was estimated by SeqFF^54^ from depth coverage calculated from cram. Single nucleotide variants (SNVs) were detected and the corresponding mutant allele frequencies (MuAFs) were estimated by BaseVar^18^ from the cram files of the discovery cohort. Based on a suggestion from the author of BaseVar, we set 100 counts into a batch and set a no-overlap sliding window with 2 million bases to detect SNVs. SNVs met any of the three filter conditions were excluded form subsequent analysis: (1) quality < 100; (2) minor allele frequency (MiAF) < 1%; (3) read depth < 500 or > 5000. Variant Effect Predictor,^63^ gnomAD,^64^ ChinaMAP,^65^ and dbNSFP^66^ database were used to annotate SNVs for obtaining useful information, including gene symbol, variant consequence, RSID, and MuAF.

### Genotype imputation

The individual genotypes of SNPs in the discovery cohort were imputed by STITCH.^20^ A reference panel of the Chinese population of the 1000 genomes project^67^ was used to initialize the ancestral haplotypes in an expectation–maximization algorithm of STITCH. We set 10 haplotypes in 100 generations as described in our previous study^18^ and imputed the SNPs in 5 million bases window with 250 kilobases buffer size. The imputed SNPs met any of the following indicators were excluded from subsequent association study: (1) *P*-value of Hardy-Weinberg balance test < 1×10^-5^; (2) MiAF < 1%; (3) the rate of genotyped individuals < 90%. (4) *I_A_*, < 0.4. *I_A_*, was defined in IMPUTE2^68^ as an information metric for estimating the accuracy of imputation. *I_A_* takes values between 0 and 1, where a value near 1 indicates that a SNP has been imputed with high certainty.

### Principal component analysis

Principal component analysis was performed by propca^69^ on the imputed genotypes of SNPs with MiAF > 5%. To decrease the memory and time of calculation, we pick up the first SNP from every 50 SNPs in sort VCF file and obtained a total of 100,000 SNPs for PCA. The genotype of wild-type, heterogeneous, and homogeneous were re-coded to 0, 1, and 2 correspondingly and stored in a modified EigenStrat^70^ format. The genetic structure of participants was displayed by the top two principal components (PCs). Linear regression was used to investigate the association between the proportion of fetal cfDNA and the top three PCs.

### Association study

In the pre-GWAS, as the individual genotypes were imputed from a pool of maternal and fetal cfDNA and the fetal cfDNA was shown to contribute to the second PC of genotype matrix, it is necessary to confirm any bias of imputed genotype for subsequent GWAS of phenotypes. A linear regression model was used to estimate the association between the proportion of fetal cfDNA and neonatal metabolism. *P*-values of multi-test were adjusted by Bonferroni method. No statistical evidence was observed for an association between fetal cfDNA and neonatal metabolism.

In the GWAS, a linear regression model was used to estimate the significance of the association between SNPs and each metabolic component by glm function in R-program, respectively. The neonatal weight/gender, maternal age/height/BMI, top three PCs of PCA were used as covariates in the regression model. *P*-values at a threshold < 5 × 10^-7^ were used for determining genome-wide significance^71^ in the discovery cohort. To validate the associations, the same method was used to calculate the significance of SNPs in the replication cohort. The statistical summary (β, *P*-value, and sample size) of association in the discovery and replication cohort were used for meta-analysis by METAL.^72^ SNPs with all *P*-values of discovery, replication, and meta-analysis pass through the threshold of < 5 × 10^-7^, < 2.68 × 10^-5^, and < 6.67 × 10^-10^ were considered to make a significant contribution to the variation of metabolite concentration. SNP with the smallest *P*-value in a gene or in one million bases window of the intergenic region was defined as the tag SNP. Besides neonatal metabolic components, we also included two maternal traits—maternal height and BMI—for GWAS. The top three PCs of PCA and maternal age were used as covariates in linear regression model. The gestational weeks and the proportion of fetal cfDNA were used as additional covariates for maternal BMI. Significant thresholds of *P*-value (*P*_discovery_ < 5 × 10^-7^, *P_r_*_eplication_ < 2.68 × 10^-5^, *P*_meta-analysis_ < 5 × 10^-10^) followed the criticizes described above.

### Trend test of allele frequency and ranked phenotypic groups

Because the association is estimated from neonatal metabolism and maternal genotype, which is un-observed in neonates in our study, we designed a trend test to estimate the association between ranked phenotypic groups and corresponding allele frequency. Firstly, we sorted the neonatal metabolite concentrations and classified the neonates into five ranked groups. Subsequently, we estimated the allele frequency for each neonatal group. Finally, we performed a trend test via prop.trend.test R function on the allele frequencies of five ranked groups.

The allele frequency of each neonatal group in the second step was estimated by:

We estimated the allele frequency in the un-observed paternal population (*p_father_*) from the allele frequency in our maternal population (*p_mother_*), male population (*p_XY_*) and female population (*p_XX_*) of gnomAD database:

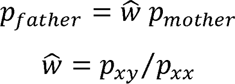

The proportions of genotypes (AA, Aa, and aa) in the un-observed paternal population were calculated according to H-W balance:

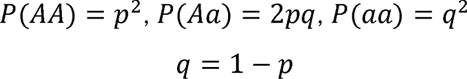

We assumed the mate-pairs of father and mother are random and they translate their allele to the offspring independently, the allele translation probability from one parent was estimated by:

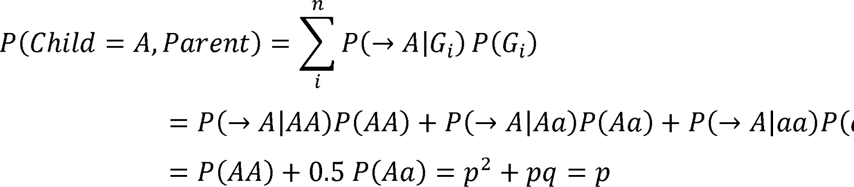

Where *P*(→ *A*\*AA*) = 1, *P*(→ *A*\*Aa*) = 0.5, *P*(→ *A*\*aa*) = 0

The proportion of neonatal genotypes (*AA, aa*, and *Aa*) were calculated by:

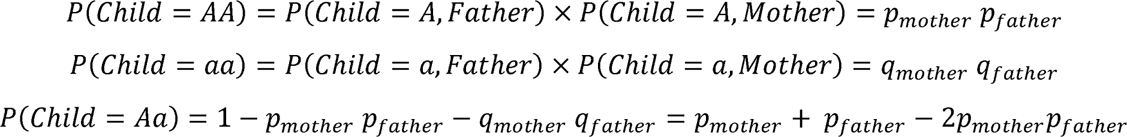

The allele frequency in neonatal population was calculated by:

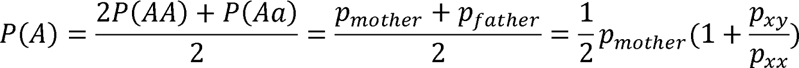

### Variant-variant interaction

SNPs with *P*-value at a threshold of < 0.01 in genome-wide association were used to study the interaction of pairwise SNPs. We set an interaction of two SNPs in linear model. *P*-value thresholds (*P*_discovery_ < 5 × 10^-7^, *P*_replication_ < 5 × 10^-7^) were used to identify the significant interaction of pairwise SNPs.

### Enrichment analysis

KEGG pathway enrichment analysis was performed in the KOBAS database.^73^ *P*-values were calculated by Fisher’s exact test and adjusted by the false discovery rate (FDR) method. SNPsea^74^ was used to screen the tissues and cells that are likely to be influenced by metabolism-associated loci. We employed 533 organs/tissues/cell types/cell lines from FANTOM^75^ for screening. Empirical P-values were calculated by permutation from null SNP sets.^76^ Cell type expression enrichment analysis was performed in large-scale single-cell transcriptome datasets of liver,^37,77^ heart,^78^ kidney,^79^ and gut^80^ via EWCE^81^ method described in our previous study.^82,83^ *P*-values were adjusted by the FDR method. Adjusted *P*-value at a threshold of < 0.05 was used to indicate the significant enrichment.

### Heritability and polygenic risk score

Kinship matrix of all participants was calculated from individual genotypes and used for heritability estimation in GCTA GREML^46^ model:

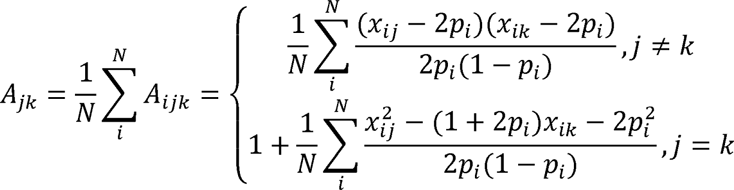

*A* refers to the kinship matrix, *j* and *k* refers to the *j*th and *k*th individual, *i* refers to the *i*th SNP of the *N* SNPs used for calculation, *x* refer to the expect dosage of genotype, *p* refers to the allele frequency of SNP. This formula indicate that we can used the kinship of mother to instead the kinship of neonates for neonatal heritability estimation. To demonstrate our hypothesis, we used the accuracy genotype from the high-depth WGS of 602 trios and showed that the pairwise correlation in neonatal kinship were highly correlated with that in maternal kinship. SNPs with *P*-value at a threshold of < 0.01 were used to estimate the heritability of neonatal metabolism and maternal height/BMI. Genotypes of SNPs were transformed to plink bed format and the kinship matrix of participants was calculated. Heritability was estimated by GREML analysis of GCTA with neonatal weight/gender, maternal age/BMI, and the top three PCs as covariates. To predict the neonatal metabolites concentration and maternal height/BMI from individual genotypes, SNPs with *P*-value at a threshold of < 0.001, < 0.005, < 0.01 and corresponding regression coefficient were used to calculate the polygenic risk score (PRS) by PRSice.^84^ A linear regression model with maternal height/BMI as covariates was used to estimate the association between PRS and phenotype. A training set of 5,000 participants in the discovery cohort was randomly selected to estimate the best-fit regression model. The remaining 3,744 participants in the discovery cohort were used to test the performance of the best-fit regression mode. Mean-square error (MSE) was used to estimate the accuracy of PRS model. The variance (VAR) was calculated from the observation. MSE/VAR was used to compare the accuracy between different metabolic components and a threshold of MSE/VAR≤0.2 was set to indicate the high-accuracy model.

## Data and code availability

The GWAS summary data sharing will be released at The National Genomics Data Center (*BF2023071213814). The necessary code of bioinformatic analysis and statistical analysis were released at GitHub (https://github.com/liuhankui/NIPT).

## Declarations

## Acknowledgments

We thank the participants for their involvement in this study. This study was supported by the grants of Primary Research & Development Plan of Jiangsu Province (BE2022736), Jiangsu Maternal and Children health care key discipline (FXK202142), Jiangsu Provincial Medical Key Discipline (Laboratory) Cultivation Unit (JSDW202214) to Prof. Ting Wang, Shenzhen Municipal of Government of China (JCYJ20180507183615145), Key research and development project of Hebei Province (Grant/Award Number 21377720D) to Prof. Jianguo Zhang.

## Author contributions

Conceptualization: Quanze He, Hankui Liu

Methodology: Hankui Liu, Quanze He

Formal analysis: Hankui Liu, Quanze He, Lu Lu, Xiaojuan Wu, Ying Xue, Chunhua Zhang, Yuxing He, Kangrong Zhao

Statistical analysis: Hankui Liu, Quanze He, Qin Zhang, Xiaojuan Wu

Interpretation: Hankui Liu, Quanze He, Jianguo Zhang, Xin Jin

Resources: Ting wang, Jianguo Zhang, Lijian Zhao, Qi wang, Benjing Wang, Huanhuan Peng, Hong Li

Funding acquisition: Ting wang, Jianguo Zhang

Supervision: Ting wang, Jianguo Zhang, Lijian Zhao, Xin Jin

Project administrator: Ting wang, Lijian Zhao, Xin Jin

Writing – original draft: Quanze He, Hankui Liu

Writing – review & editing: Quanze He, Hankui Liu, Ting wang, Jianguo Zhang, Lijian Zhao, Xin Jin, Liping Guan, Lu Lu, Jun Mao, Xiangwen Peng

All authors read and approved the final manuscript.

---

Figure S1. The distribution of 75 metabolic components and maternal height/BMI.

Figure S2. The quality control metric of sequencing data and SNVs.

Figure S3. The proportion of fetal cfDNA was estimated from 1% to 30% with a median of 10%. The proportion of fetal cfDNA was significantly affected by the maternal weight and have an impact on the PC2 of imputation genotype (A). The maternal age (mAge), maternal height (mHeight), maternal BMI (mBMI), neonatal weight (nWeight) and neonatal sex (nSex) were shown to significantly affected the neonatal metabolism. In contrast, there are no statistical evidence for a significant association of neonatal metabolism and the maternal ancestry or the proportion of fetal cfDNA. Red horizontal line refers to the significant threshold (0.05/[7*75]) (B).

Figure S4. The Q-Q plots of the associations of 75 neonatal metabolic components and maternal height/BMI. λ_gc_ refers to the genomic inflation factor calculated from the median *P*-value.

Figure S5. The correlation of maternal allele frequency and neonatal/parental allele frequency (A), estimated allele frequency and observed allele frequency (B), maternal kinship and neonatal kinship (C) in 602 trios with high-depth WGS data.

Figure S6. The accuracy of PRS model of neonatal metabolic components.

## Key resource table

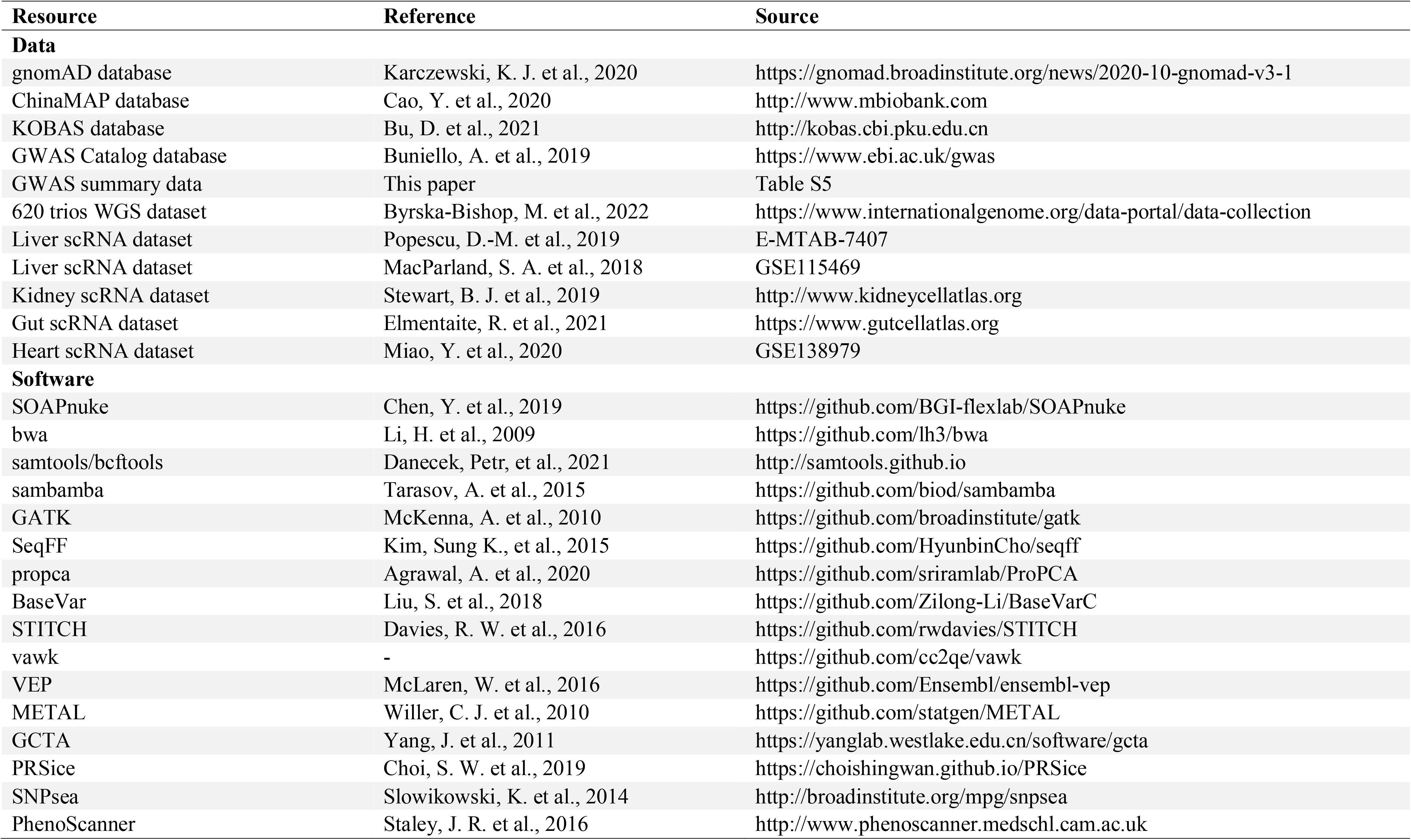
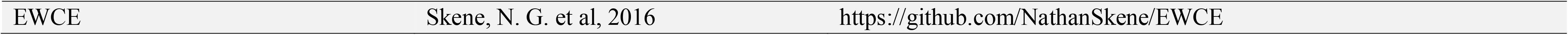

